# The Phosphate-Specific Transport System Gene *pstA1* Contributes to Rifampin Tolerance in *Mycobacterium tuberculosis*

**DOI:** 10.1101/2025.10.31.685634

**Authors:** Carina Danchik, G. V. Anoushka Chinmayi, Gi Yong Lee, Joel S. Bader, Hyungjin Eoh, Petros C. Karakousis

## Abstract

Tuberculosis (TB) caused an estimated 10.8 million new cases and 1.25 million deaths in 2023. Antibiotic tolerance, the ability of bacteria to survive exposure to bactericidal antibiotics without genetic resistance mutations, contributes to the need for prolonged treatment. Targeting antibiotic tolerance mechanisms could promote accelerated clearance of *Mycobacterium tuberculosis* (*Mtb*), thereby improving medical adherence. Our previous forward genetic screen for rifampin tolerance genes identified *pstA1*, which is involved in phosphate-specific import. The ABC-type transporter permease PstA1 has previously been implicated in *Mtb* virulence, as mutants lacking this gene exhibit defective survival during phosphate limitation *in vitro*, in infected macrophages, and in immunocompetent mice. The minimum inhibitory concentration (MIC) of rifampin was not altered in a *pstA1* deletion mutant (Δ*pstA1*), suggesting this difference in susceptibility is not due to antibiotic resistance. Consistent with a role in rifampin tolerance, time-kill assays revealed a shift in the mean duration of killing (2-log reduction) from 1.6 days in wild-type to 1.0 day in Δ*pstA1*, and complementation partially restored this phenotype. We found that *pstA1* is specifically required for *Mtb* survival in the absence of exogenous inorganic phosphate and is important for adaptation to growth in culture without detergent, and within macrophages in an interferon-γ-dependent manner. Differential expression analysis revealed that Δ*pstA1* exhibited substantial transcriptional reprogramming with 58 differentially expressed genes, including altered expression of metabolic, DNA damage repair, and secretory pathways. PstA1 represents a novel drug target, and inhibitors could serve as adjunctive therapies to shorten treatment times, reducing opportunities for drug resistance emergence.

**Importance:** Tuberculosis remains one of the world’s deadliest infectious diseases, causing over a million deaths annually. Current treatment requires months of antibiotic therapy, and poor adherence to these lengthy regimens contributes to the emergence of drug-resistant strains that are increasingly difficult to treat. A major barrier to shorter treatment is antibiotic tolerance, which allows bacteria to survive drug exposure without genetic resistance mutations. This study identifies the phosphate transport component PstA1 as a critical factor enabling *Mycobacterium tuberculosis* to tolerate the frontline antibiotic rifampin. Bacteria lacking *pstA1* are eliminated more rapidly by rifampin, demonstrating that this transporter actively promotes bacterial survival during treatment. These findings suggest that drugs targeting PstA1 could be combined with standard antibiotics to accelerate bacterial clearance, potentially shortening treatment duration and improving patient adherence. Such adjunctive therapies targeting tolerance mechanisms represent a promising strategy to combat tuberculosis and reduce the global burden of drug-resistant disease.

## Introduction

Tuberculosis (TB) remains the leading cause of death from a single infectious agent, causing an estimated 10.8 million new cases and 1.25 million deaths in 2023 alone [1]. While effective regimens exist and three new drugs have been approved to treat TB since 2010, additional advances are needed to shorten treatment duration and improve treatment completion [2]. Current first-line TB treatment requires the use of multiple antibiotics for a minimum of six months to prevent relapse. A key reason for this lengthy treatment time is the presence of residual populations of bacteria that exhibit antibiotic tolerance– a phenotypic and reversible state in which bacteria become transiently insusceptible to antibiotics, requiring longer exposures to achieve bacterial eradication [3]. Bacteria can use a variety of adaptive mechanisms, collectively termed phenotypic tolerance, to adapt to and survive under stress conditions experienced within the host as well as antibiotic pressure [4]. The molecular mechanisms underlying antibiotic tolerance in *Mtb* are complex, involving transcriptional stress responses, metabolic adaptations, activation of drug efflux pumps, and changes in cell wall permeability [5]. Direct disruption of *Mtb* antibiotic tolerance mechanisms is expected to result in more rapid bacillary killing, leading to shorter treatment times that may improve medical adherence and decrease opportunities for the development of antibiotic resistance [6].

We previously used transposon mutant sequencing (Tn-seq) to identify *Mtb* genes required for survival during exposure to bactericidal concentrations of the first-line drug rifampin. Among the top hits from this unbiased genome-wide approach were the phosphate-specific transport (Pst) genes *rv0820/phoT*, *rv0929/pstC2*, and *rv0930/pstA1* [7]. Notably, their homologs were also identified in our similar forward genetic screen in which *M. avium* was exposed to rifabutin, providing additional confidence in their significance and suggesting that Pst genes contribute to rifamycin tolerance across mycobacterial species [6]. Pst systems are comprised of four essential components: PstA and PstC are permeases that form a transmembrane channel complex; PstS is an extracellular protein that senses and captures phosphate from the environment; and PstB is an intracellular ATPase that provides energy for active transport of phosphate against its concentration gradient [8–10]. PstA1, which is co-expressed as part of the putative *Mtb* operon *pstS3-pstC2-pstA1* [11], is an important virulence factor required for *Mtb* survival during acidic stress [12] and phosphate depletion [13, 14], as well as for *Mtb* survival within macrophages [15] and for resistance to IFN-γ dependent immunity in mice [14]. PstA1 also plays important regulatory roles, as several substrates of the ESX-5 secretion system, as well as membrane vesicles, are hypersecreted in Δ*pstA1* mutants [16, 17]. Each of these phenotypes is mediated at least in part through the constitutive activation of RegX3, a response regulator of the PhoPR two-component system, in the absence of PstA1 [18, 19]. This regulatory connection links phosphate sensing to broader stress responses and virulence factor expression. Previous studies have shown that a *pstA1* deletion mutant was more susceptible to several drug combinations *in vitro* and to rifampin in a mouse model [20], providing initial evidence for a role in antibiotic tolerance. However, the molecular mechanisms underlying this phenotype have not been fully elucidated.

In this study, we have comprehensively characterized the role of *pstA1* in rifampin tolerance using phenotypic assays, transcriptomics, and metabolomics to explore the underlying molecular mechanisms of its increased susceptibility. We have also studied the role of PstA1 in *Mtb* growth and survival in phosphate-rich and phosphate-depleted models of nutrient starvation and within macrophages, providing insights into how phosphate transport contributes to bacterial adaptation under diverse stress conditions. Our findings reveal new mechanistic insights into antibiotic tolerance and highlight the potential for targeting phosphate transport pathways in TB treatment.

## Results

### Phosphate transport gene deletions confer rifampin hypersusceptibility

Our Tn-seq screen to identify *Mtb* genes required for tolerance to the sterilizing drug rifampin identified three Pst genes, *rv0820/phoT*, *rv0929/pstC2*, and *rv0930/pstA1*, among the top hits [7]. Their homologs were also found in a similar Tn-seq study of *M. avium* exposed to rifabutin [6]. Notably, *pstC2* and *pstA1* are downstream components of an *Mtb* operon including *pstS3* [22, 37]. To confirm that these three genes are co-transcribed in the H37Rv strain under our experimental conditions, we performed RT-PCR analysis using primers spanning the intergenic and intragenic regions of this operon. Our results confirmed that *pstS3*, *pstC2*, and *pstA1* are co-expressed as a single polycistronic mRNA transcript (Fig 1), validating the operonic structure for subsequent studies.

**Figure 1.**
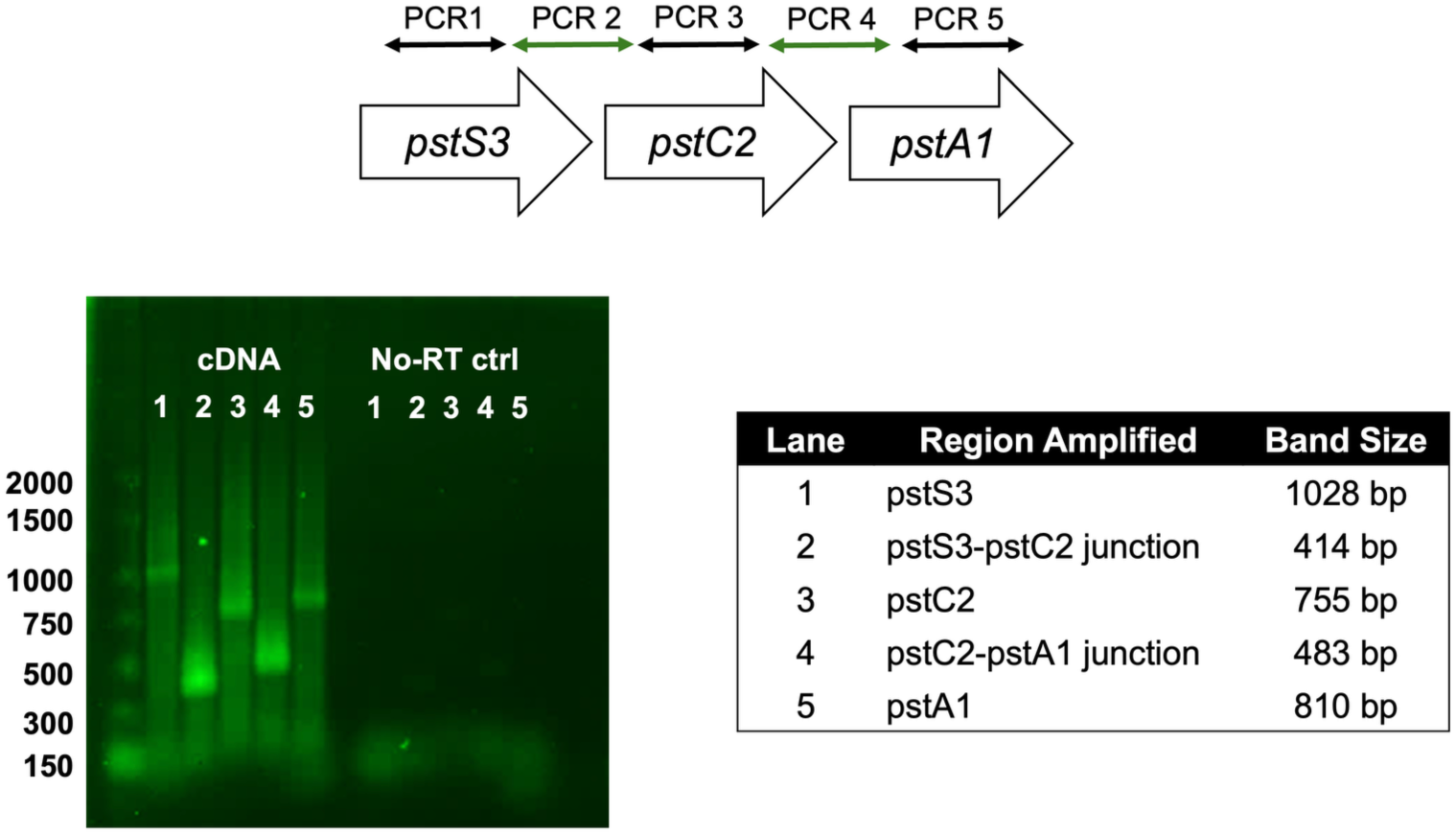
The *pstS3-pstC2-pstA1* genes are co-transcribed as an operon. RT-PCR analysis of H37Rv total RNA using primers spanning intragenic regions within *pstS3* (lane 1), *pstC2* (lane 3), and *pstA1* (lane 5), or intergenic junctions between *pstS3-pstC2* (lane 2) and *pstC2-pstA1* (lane 4). Amplification of all regions from cDNA confirms continuous transcription across the three genes. No-RT controls (right panel) show the absence of genomic DNA contamination. Expected amplicon sizes are indicated in the table.

Next, we generated individual targeted deletion strains for each of these three genes using the ORBIT cloning system [21]. Cloning was validated via PCR of the 5’ and 3’ ends of the insertion site, as well as with whole genome sequencing for Δ*pstA1* (SeqCenter, Pittsburgh, PA) (Supplementary Figure S1). To validate our Tn-seq findings, we assessed the activity of rifampin against each deletion strain using standard time-kill assays following addition of 2.0 μg/ml rifampin. All three deletion strains showed increased susceptibility to rifampin treatment compared to wildtype controls harboring empty vectors. At day 6, Δ*phoT* showed a non-significant 0.9 log₁₀ reduction in CFU compared to control (adjusted P = 0.358; Fig. 2A). Δ*pstC2* demonstrated a more pronounced phenotype with an additional 1.3 log₁₀ reduction (adjusted P = 0.000189, Fig. 2B). Δ*pstA1* exhibited the strongest susceptibility to rifampin, showing an additional 1.9 log₁₀ reduction in viable bacteria (adjusted P = 0.00742) relative to the isogenic wild-type control (Fig. 2C). The minimum duration of killing (MDK), defined as the time required to achieve a 2-log₁₀ reduction from the starting bacterial density, decreased by 0.4 days for Δ*phoT*, 1.1 days for Δ*pstC2*, and 0.6 days for Δ*pstA1* relative to controls. Consistent with a role for each of these genes in *Mtb* tolerance, rather than resistance, to rifampin, each deletion mutant showed minimal alterations in rifampin MIC values. The MIC values of rifampin against Δ*phoT* and Δ*pstC2* were 0.03 μg/ml and that of Δ*pstA1* was 0.016 μg/ml (wild type, 0.03-0.06 μg/ml).

**Figure 2.**
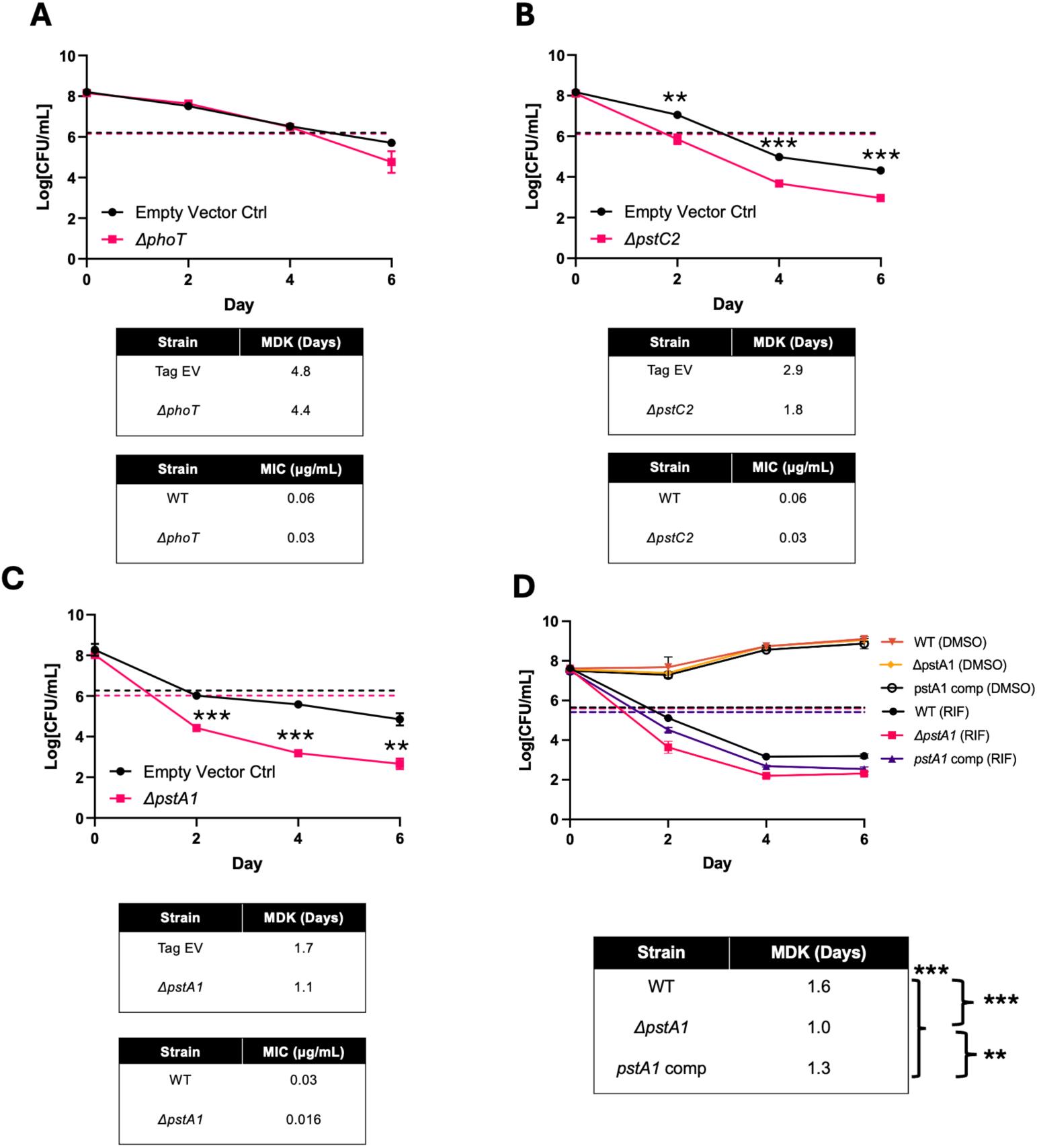
Deletion of phosphate-specific transport genes enhances rifampin susceptibility. Time-kill assays were performed in 7H9 medium with 2 μg/mL rifampin for (A) ΔphoT, (B) ΔpstC2, and (C) ΔpstA1 in parallel with an empty vector control. (D) Time-kill assays performed with 7H9 medium with 0.5 μg/mL rifampin on wild-type, ΔpstA1 and pstA1 complement strains. Dotted lines indicate 2-log₁₀ reduction from starting inoculum. Tables show minimum duration of killing (MDK) to achieve 2-log₁₀ reduction and rifampin MIC values for each strain. Statistical significance for CFU comparisons at individual timepoints (A-C) was determined by unpaired t-test with Holm-Šídák correction for multiple comparisons. Statistical significance for MDK values (D) was determined by one-way ANOVA with Tukey’s multiple comparisons test (*P ≤ 0.05; **P ≤ 0.01; ***P ≤ 0.001).

### Complementation of *pstA1* partially restores wild-type rifampin tolerance

Given the robust phenotype observed with Δ*pstA1*, and the possibility that deletion of *pstC2* could have polar effects on *pstA1*, our subsequent studies focused on Δ*pstA1*. To validate the *pstA1* deletion phenotype, we constructed a genetic complementation strain (Δ*pstA1::pstA1*). Since *pstS3*, *pstC2*, and *pstA1* are co-transcribed as an operon, we designed our complementation construct to include appropriate regulatory elements. The *pstA1* coding sequence fused to the putative operonic promoter region (422 bp upstream of *pstS3* plus the first 75 bp of *pstS3*) was cloned into the episomal mycobacterial vector pMH94 and introduced into the Δ*pstA1* strain. Complementation was confirmed by PCR of insert junctions and whole genome sequencing (Supplementary Figure S1). Time-kill assays revealed that the minimum duration of killing (MDK) of rifampin for the complement strain was 1.3 days, as compared to 1.6 days for the wild type and 1.0 day for ΔpstA1 (Fig. 2D). MDK values differed significantly between all three strains (WT vs ΔpstA1: p = 0.00004; WT vs pstA1 comp: p = 0.004; ΔpstA1 vs pstA1 comp: p = 0.0011, one-way ANOVA with Tukey’s multiple comparisons test). Thus, genetic complementation of Δ*pstA1* partially restored wild-type rifampin tolerance.

### Δ*pstA1* exhibits phosphate-specific survival defects while maintaining rifampin susceptibility under starvation conditions

To characterize the broader physiological role of *pstA1*, we examined bacterial survival under various stress conditions relevant to mycobacterial pathogenesis and persistence. As multiple forms of nutrient starvation are known to induce a tolerance phenotype [13, 22, 23], we tested the ability of Δ*pstA1* to survive in both the PBS nutrient starvation model used in our prior screen [7] and during inorganic phosphate limitation [13].

During nutrient starvation in phosphate-buffered saline containing the detergent tween-80, both wild-type and Δ*pstA1* strains showed equivalent survival over a 14-day adaptation period, with no significant differences observed between strains (Fig. 3A). *ΔpstA1* retained enhanced rifampin susceptibility under nutrient-starved conditions. When rifampin (2.0 μg/ml) was added after the 14-day starvation period, the deletion mutant showed significantly enhanced killing compared to wild-type, with additional reductions in viability becoming apparent by day 2 (1.4 log₁₀ reduction; adjusted P = 0.000184) and persisting through day 6 (1.6 log₁₀ reduction; adjusted P = 0.000184) (Fig. 3B).

**Figure 3.**
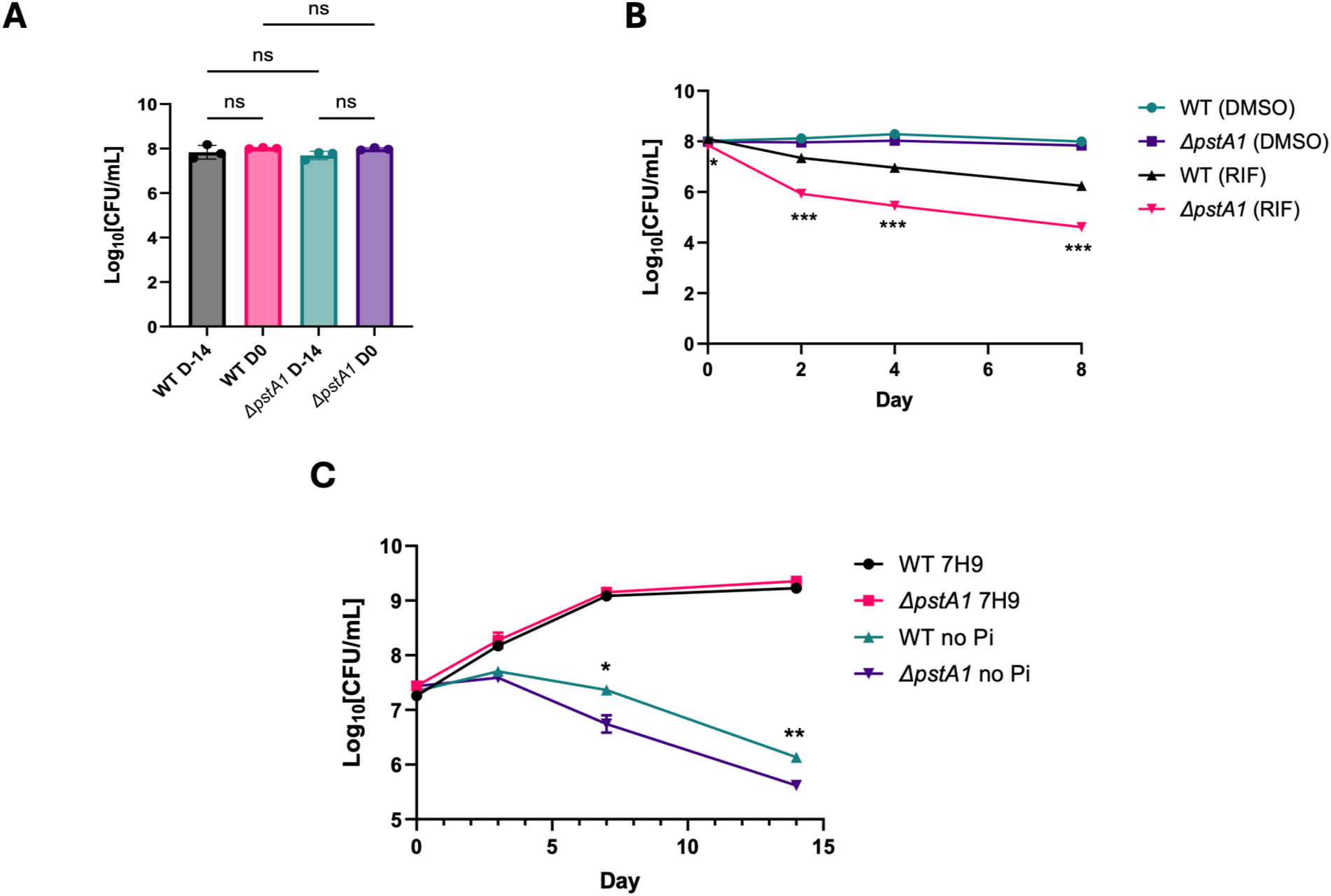
Δ*pstA1* survival under nutrient starvation and phosphate depletion. (**A**) Wild-type H37Rv and Δ*pstA1* survival after exposure to PBS-Tween for 14 days. No significant differences were observed by one-way ANOVA followed by Tukey’s multiple comparisons test. (**B**) Δ*pstA1* maintained increased rifampin sensitivity after 2 weeks of PBS pre-treatment. Bacteria were treated with 2 μg/mL rifampin or DMSO vehicle control. Statistical differences indicated are for WT (RIF) vs Δ*pstA1* (RIF). Representative of two independent experiments. (**C**) Δ*pstA1* shows no growth defect in phosphate-rich 7H9 medium but exhibits accelerated death under phosphate depletion (no Pi). Statistical differences shown are between WT no Pi and Δ*pstA1* no Pi conditions. Representative of three independent experiments.

Since phosphate depletion is known to induce the stringent response, a key antibiotic tolerance pathway in *Mtb* [24, 25], and the *pstS3-pstC2-pstA1* operon and *phoT* are upregulated under low phosphate conditions [13], we tested the ability of Δ*pstA1* to survive in the absence of exogenous inorganic phosphate. Phosphate-specific depletion revealed a clear survival disadvantage for Δ*pstA1*. While both wild-type and Δ*pstA1* strains exhibited similar growth and survival characteristics in standard phosphate-rich 7H9 broth, Δ*pstA1* showed significantly reduced viability under phosphate-depleted conditions. In modified 7H9 broth lacking inorganic phosphate, the deletion strain demonstrated an additional survival defect with a 0.7 log₁₀/ml decrease by day 7 (adjusted P = 0.0115) and a 0.6 log₁₀/ml decrease by day 14 (adjusted P = 0.00181) relative to wild type (Fig. 3C). These results confirm the role of *pstA1* in phosphate acquisition under limiting conditions and demonstrate its requirement for growth in low-phosphate environments.

### Δ*pstA1* exhibits survival defects in IFN-γ-activated macrophages

To assess the relevance of our findings to mycobacterial pathogenesis, we next examined the role of *pstA1* in macrophage infection and intracellular survival as a model of the natural host environment. Macrophages represent the primary host cell niche for *Mtb* and create a hostile intracellular environment characterized by nutrient limitation, acidic pH, and antimicrobial factor exposure [15]. We infected PMA-differentiated THP-1 macrophages with wild-type and Δ*pstA1* strains at a multiplicity of infection (MOI) of 1:1 and monitored intracellular bacterial survival over 4 days. In resting macrophages, there was no significant difference in bacterial burden between wild-type and Δ*pstA1* at any time point measured (days 0, 2, and 4 post-infection; Fig. 4A), suggesting that *pstA1* is not required for initial invasion or survival in resting macrophages. However, when macrophages were pre-activated with IFN-γ, a key cytokine that enhances macrophage antimicrobial functions, Δ*pstA1* showed a significant survival defect that became pronounced by day 4 post-infection. While wild-type bacteria maintained relatively high intracellular numbers (4.2×10⁶ CFU/ml), the Δ*pstA1* strain showed reduced survival (2.43×10⁶ CFU/ml), representing a statistically significant difference (adjusted P = 0.0098; Fig. 4B). This result confirms previous reports that *pstA1* is specifically required for resistance to IFN-γ-mediated host immunity and suggests a role in counteracting activated macrophage antimicrobial mechanisms [15].

**Figure 4.**
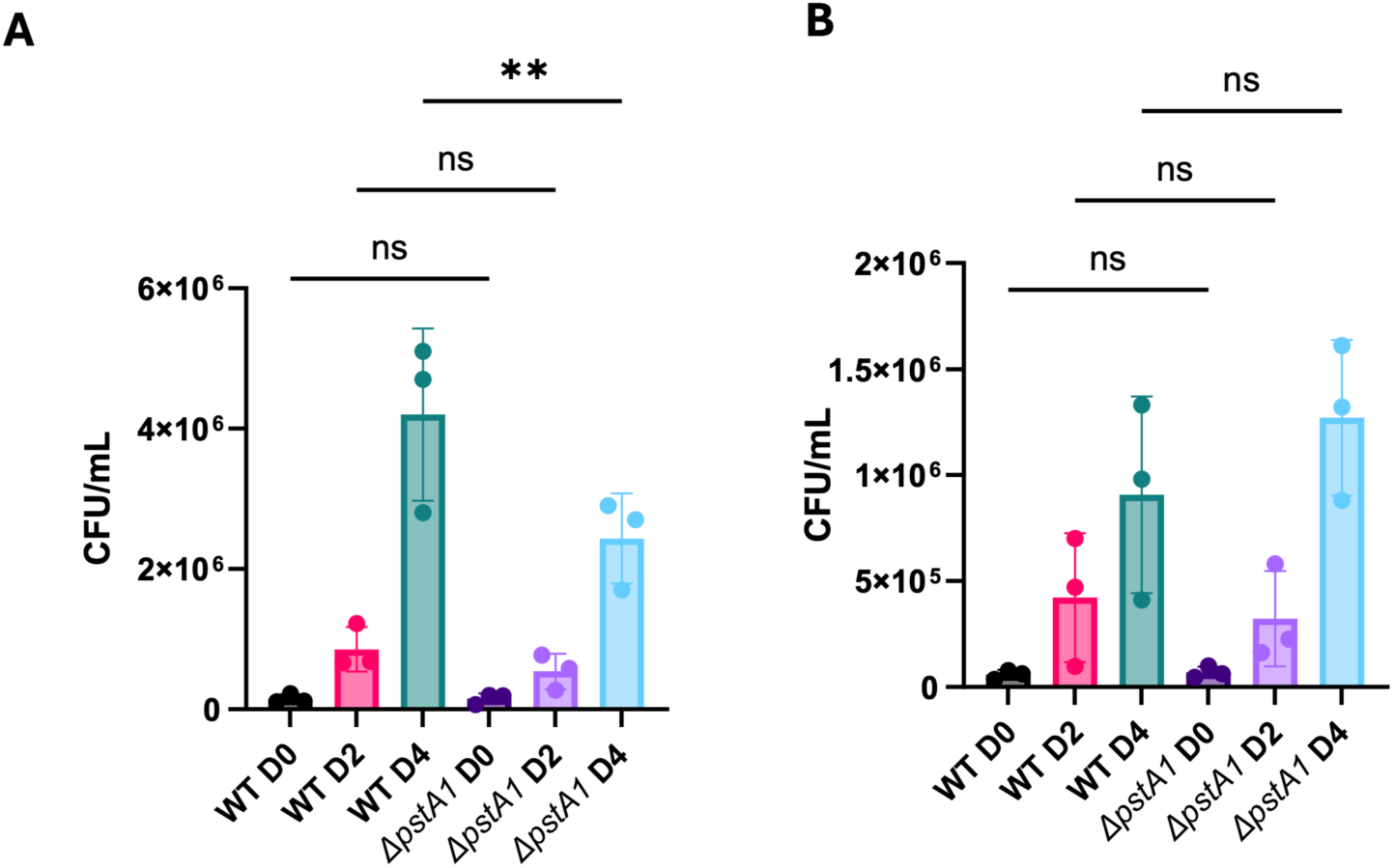
Δ*pstA1* exhibits an IFN-γ-dependent growth defect within macrophages. Bacterial burden within PMA-differentiated THP1 cells after 4 hours of infection (D0) or 2 or 4 days without (A) or with (B) 24 hr IFN-γ polarization before infection. One way ANOVA was performed followed by Sidak’s multiple comparisons test to determine significance. Adjusted p-value ≤0.05 = *; ≤0.01 = **; ≤0.001 = ***.

### The contribution of PstA1 to *Mtb* rifampin tolerance is not mediated through the accumulation of inorganic polyphosphate

Having established that Δ*pstA1* exhibits enhanced rifampin susceptibility, we next sought to investigate the underlying mechanisms responsible for this phenotype. Induction of the *Mtb* stringent response and accumulation of the multifunctional stress signaling molecule, inorganic polyphosphate (polyP), has been linked to antibiotic tolerance in *Mtb* [26–30]. Since RegX3, a two-component response regulator that controls the *Mtb* phosphate starvation and stringent responses [13, 31], is constitutively activated upon *pstA1* deletion [14], we measured intracellular polyP levels as a proxy for stringent response activation. No significant differences in polyP accumulation were observed between wild-type and Δ*pstA1* strains across different growth phases (Fig. 5A). Wild-type polyP content ranged from 3.072 to 5.536 ng/μg total protein, while Δ*pstA1* levels ranged from 3.673 to 5.915 ng/μg total protein.

**Figure 5.**
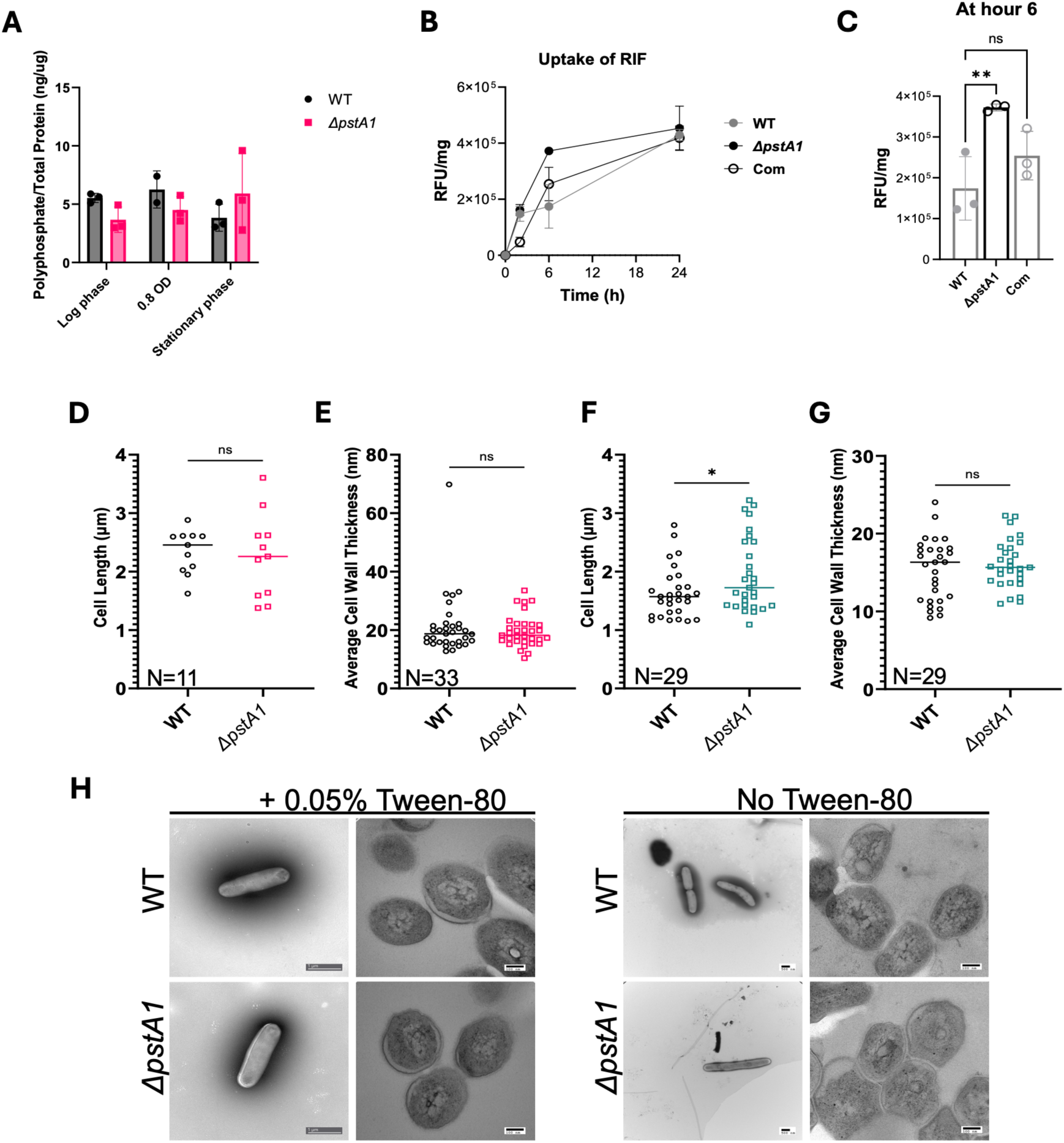
Rifampin uptake kinetics and cell morphology analysis of Δ*pstA1*. (**A**) Intracellular polyphosphate levels across different growth phases for wild-type and Δ*pstA1* normalized to total protein concentration. No significant differences were observed by one-way ANOVA followed by Tukey’s multiple comparisons test. (**B-C**) Rifampin uptake kinetics in wild-type, Δ*pstA1*, and *pstA1* complement strains measured in a filter culture system. Bacteria were treated with 1.0 μg/mL rifampin and uptake was calculated as (initial rifampin concentration in media at 0 h -rifampin concentration at each timepoint) normalized to protein concentration. (**D-E**) Cell length and average cell wall thickness of wild-type and Δ*pstA1* cultures grown in 7H9 medium with 0.05% Tween-80 during log phase. (**F-G**) Cell length and average cell wall thickness of cultures grown without Tween-80. Cell length measurements: n=11 bacteria per strain (with Tween-80), n=29 bacteria per strain (without Tween-80). Cell wall thickness was determined by averaging two measurements per bacterium: n=33 bacteria per strain (with Tween-80), n=29 bacteria per strain (without Tween-80). (**H**) Representative transmission electron microscopy images of whole bacteria or thin sections from log-phase cultures grown with or without 0.05% Tween-80. Data represent mean ± SD. Statistical significance was determined by one-way ANOVA followed by Tukey’s multiple comparisons test (**B-C**) or unpaired t-test (**D-G**). \*\**P* ≤ 0.01; \**P* ≤ 0.05; ns, not significant.

### Δ*pstA1* exhibits increased rifampin uptake

We hypothesized that *pstA1* deficiency might be associated with increased intrabacterial accumulation of rifampin. Using a previously validated device that enables timed start-stop measurements of antibiotics remaining in spent media [32], we quantified rifampin content in the wild-type and mutant strains The kinetics of rifampin uptake revealed that Δ*pstA1* displayed significantly faster and greater rifampin accumulation compared to wild-type at 6 hours post-exposure, suggesting increased uptake and/or decreased efflux of rifampin (Fig. 5B-C).

### Cell morphology analysis reveals minimal structural differences

To determine whether the observed increase in rifampin uptake was attributable to gross alterations in cell wall structure of the mutant, we examined bacterial morphology using electron microscopy. Under standard growth conditions including Tween-80 to prevent bacterial clumping, we observed no significant difference in cell length between the two strains (average bacterial length= 2.34 μm for wild type and 2.26 μm for Δ*pstA1;* Fig. 5D). Likewise, the strains showed similar average cell wall thicknesses of 21.18 nm and 19.61 nm, respectively (Fig. 5E). Substantial variations in cell wall thickness were observed for both wild-type and Δ*pstA1* which appear similar to previously reported lipid inclusions [33]. There were no significant differences in minimum, maximum, or average cell wall thickness between the two strains.

Since use of detergent can alter cell wall permeability [34], the experiment was repeated using cultures grown without Tween-80 (Fig. 5F-G). Cell length decreased for both strains in detergent-free broth, from an average of 2.34 μm to 1.64 μm for wild-type and from 2.26 μm to 1.95 μm for Δ*pstA1* (difference= 0.31 μm; P = 0.0329; Fig. 5F). Cell wall thickness did not vary notably between strains with averages of 15.37 nm for wild type and 16.06 nm for Δ*pstA1*. The large variations in cell wall thickness seen in cultures grown with Tween-80 were not present under detergent-free conditions (Fig. 5H). These results demonstrate that Δ*pstA1* exhibits increased rifampin accumulation without major changes in cellular morphology.

### Transcriptomic analysis reveals extensive metabolic and regulatory reprogramming in Δ*pstA1*

To further explore the molecular mechanisms underlying the increased rifampin susceptibility of Δ*pstA1* and identify potential cellular pathways contributing to enhanced drug uptake, we performed transcriptomic analysis using RNA sequencing. Wild-type and Δ*pstA1* strains were exposed to a sub-inhibitory concentration of rifampin (7.5 ng/ml, representing 1/4 MIC) for 30 or 60 minutes. This concentration and timing were selected to capture early transcriptional responses while minimizing the global transcriptional inhibition that occurs at bactericidal concentrations of rifampin. During axenic growth in nutrient-rich broth lacking rifampin, Δ*pstA1* exhibited substantial transcriptional reprogramming compared to wild type, with 58 differentially expressed genes (DEGs) identified: 49 upregulated and 9 downregulated (adjusted P < 0.05, |log₂ fold change| > 1; Fig. 6A). Several genes known to be part of the phosphate regulon showed significant upregulation, including *pstS3* (log₂ fold change = 3.22), *pstC2* (log₂ fold change = 2.50), and the two-component regulatory system genes *senX3* (log₂ fold change = 1.59) and *regX3* (log₂ fold change = 1.77), indicating constitutive activation of phosphate starvation signaling.

**Figure 6.**
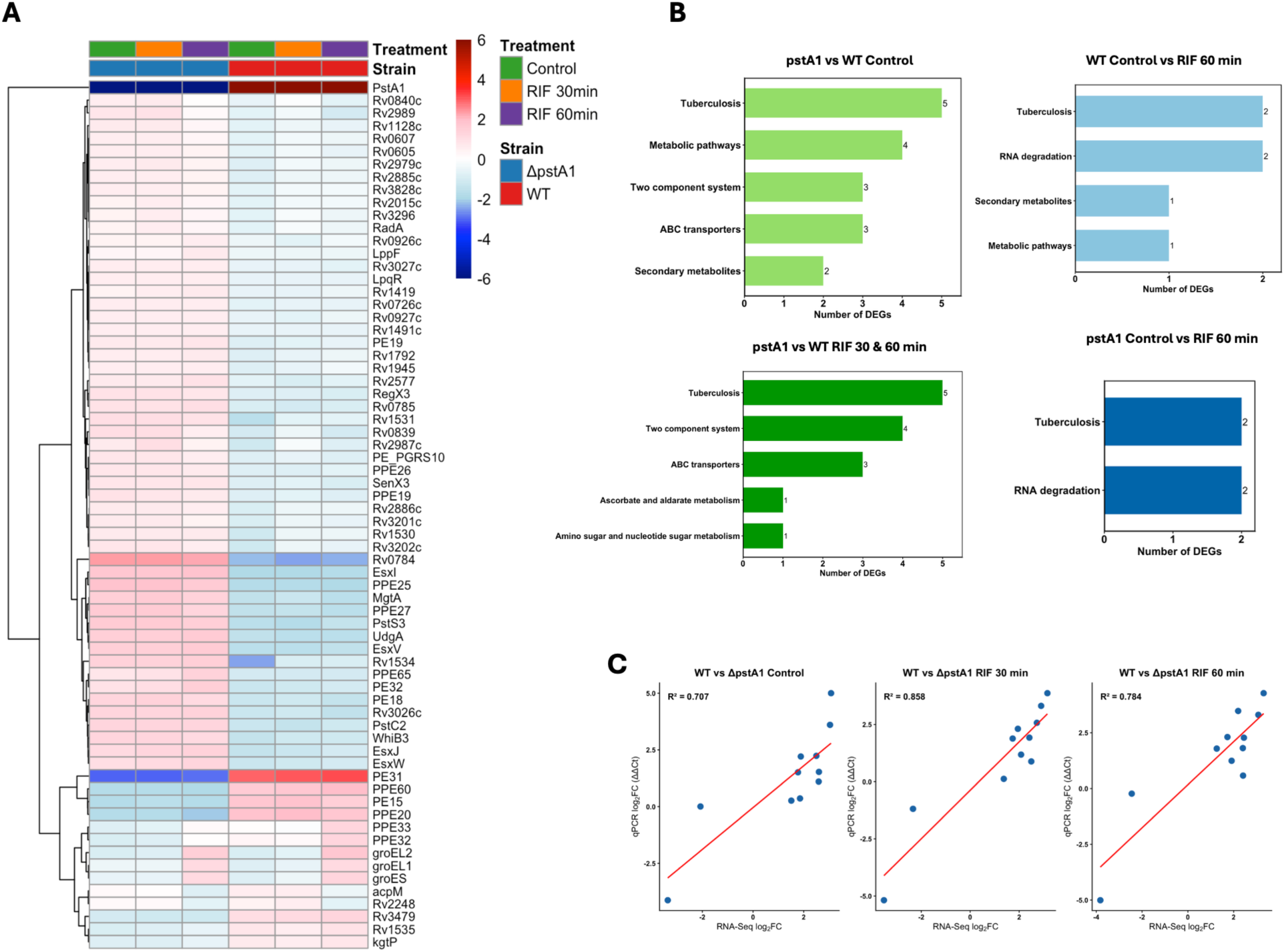
Transcriptional profiling of Δ*pstA1* with and without rifampin treatment. (**A**) Heatmap of differentially expressed genes (DEGs) in wild-type and Δ*pstA1* strains at baseline and after rifampin treatment (30 and 60 min). Genes were filtered for absolute log₂ fold change ≥ 1 and adjusted *P* < 0.05. Each row represents a gene, and each column represents a biological replicate. Color scale indicates log₂ fold change relative to wild-type control. (**B**) Functional enrichment analysis of DEGs for key comparisons: Δ*pstA1* vs wild-type control (top left), wild-type control vs wild-type with 60 min rifampin treatment (top right), Δ*pstA1* control vs Δ*pstA1* with rifampin treatment at 30 and 60 min (bottom left), and wild-type control vs Δ*pstA1* with 60 min rifampin treatment (bottom right). Bar plots show the number of DEGs in each functional category. Correlation between RNA-seq and RT-qPCR validation for selected genes across three comparisons. Each point represents a gene, with correlation coefficients (R²) indicated for each comparison.

KEGG pathway enrichment analysis revealed that genes involved in biosynthesis and metabolic pathways were prominently represented among the DEGs (Fig. 6B). The differentially expressed genes included fourteen PE/PPE family genes (9 upregulated, 5 downregulated), with upregulation of members of the ESX-5 locus and downregulation of *pe31* (log₂ fold change = −6.02) and *ppe60* (log₂ fold change = −3.37). Genes encoding four ESAT-6-like proteins were upregulated, namely *esxI* (log₂ fold change = 3.65), *esxJ* (log₂ fold change = 2.59), esxV (log₂ fold change = 3.14), and *esxW* (log₂ fold change = 2.35). The redox-responsive transcriptional regulator *whiB3* (log₂ fold change = 2.60), a key regulator of thiol metabolism and oxidative stress responses, was also significantly upregulated [12]. Additionally, eleven genes involved in DNA metabolism were upregulated in the mutant at baseline, including resolvases, transposases, and DNA helicases. Rifampin treatment led to a progressive reduction in the number of DEGs between strains. After 30 minutes of rifampin exposure, 41 genes remained differentially expressed between strains. After 60 minutes, 41 genes were differentially expressed, with 36 genes showing consistent differential expression across all three conditions (baseline, 30 min, and 60 min). A comprehensive Venn diagram analysis of DEG overlap across all treatment conditions is shown in Supplementary Figure S2. The DNA metabolism genes were no longer significantly differentially expressed following rifampin treatment, while the core set of 36 genes included the phosphate regulon genes, all four ESAT-6-like protein genes, twelve PE/PPE family genes, and *whiB3*.

RT-qPCR analysis of 11 selected genes confirmed the expression patterns observed in the RNA-seq dataset, with correlation coefficients exceeding 0.7 for all comparisons (Fig. 6D; Supplementary Figure S3). Validated genes included members of the phosphate starvation response (*pstC2*, log₂ fold change = 2.50; *regX3*, log₂ fold change = 1.77; *senX3*, log₂ fold change = 1.59), metabolic enzymes (*udgA*, log₂ fold change = 3.08; *mgtA*, log₂ fold change = 3.04), the stress response regulator *whiB3* (log₂ fold change = 2.60), and PE/PPE family protein genes (*ppe60*, log₂ fold change = −3.37; *ppe65*, log₂ fold change = 1.89). A complete list of all DEGs across all experimental conditions is provided in Supplementary Table S2.

### Metabolomic analysis reveals altered antioxidant metabolites in Δ*pstA1*

To understand the biochemical consequences of increased rifampin accumulation in the deletion mutant, we performed targeted metabolomics analysis of rifampin-treated bacteria and analysis was focused on antioxidant pathways. We analyzed the intracellular abundance of 16 metabolites involved in antioxidant biosynthesis and redox homeostasis in the Δ*pstA1* mutant, wild-type, and complement strains at 6 hours post-rifampin exposure. The heatmap analysis of all measured antioxidant metabolites revealed distinct metabolic profiles between strains (Fig. 7A). While most metabolites, including L-glutamate, glycine, NADPH, glutathionylspermidine, and various glutathione derivatives, showed trends toward increased abundance in Δ*pstA1*, these changes did not reach statistical significance. The complement strain generally displayed metabolite levels which were intermediate between wild-type and the deletion mutant, consistent with partial restoration of the metabolic phenotype.

**Figure 7.**
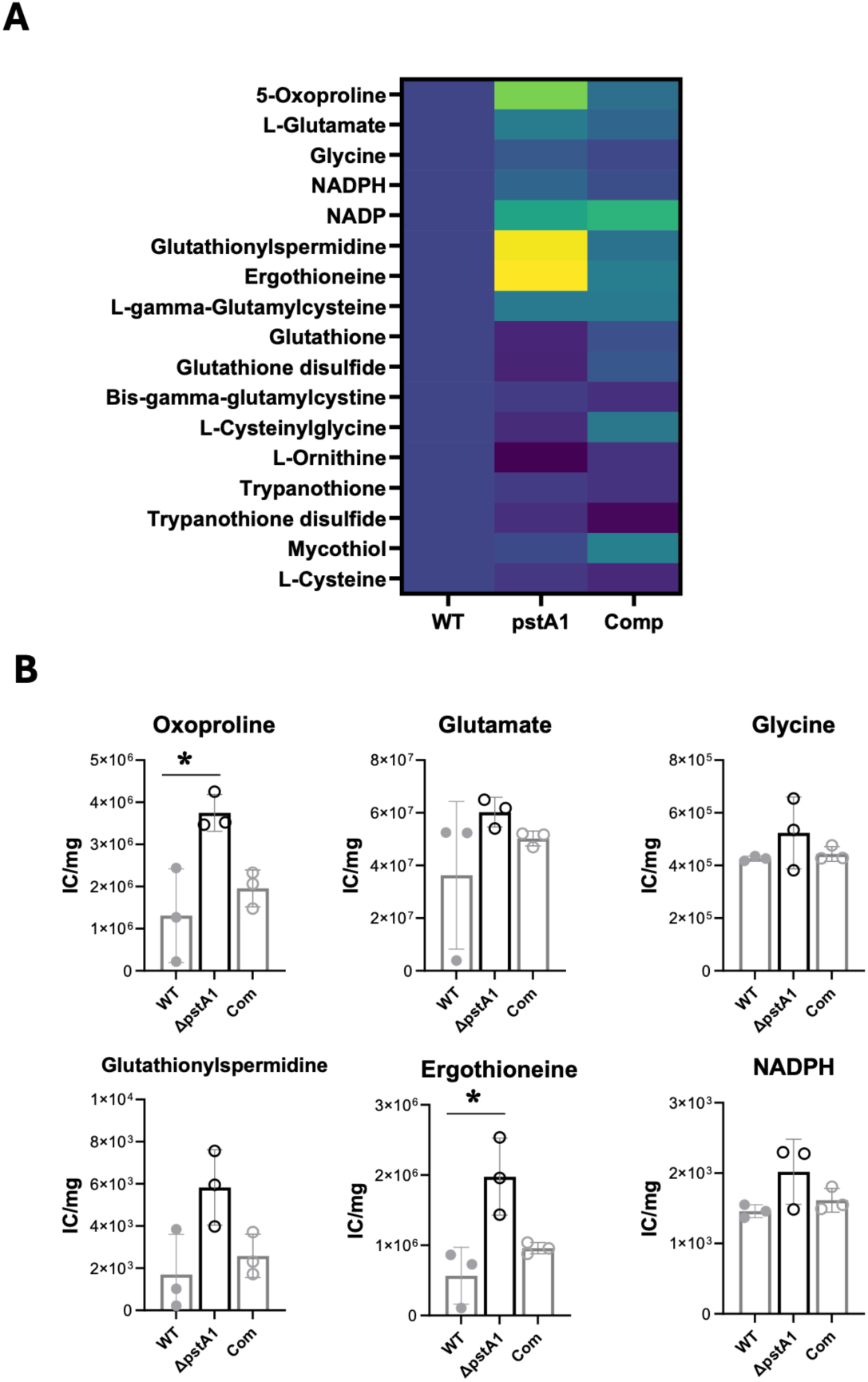
Metabolomic profiling reveals elevated antioxidant metabolites in rifampin-treated Δ*pstA1*. (**A**) Heatmap showing relative abundances of intracellular metabolites involved in antioxidant and redox pathways in wild type, Δ*pstA1*, and *pstA1* complement strains after 6 hours of rifampin exposure. Color scale indicates relative metabolite abundance (normalized to internal standards and protein concentration). (**B**) Quantification of selected antioxidant pathway metabolites. Bars represent mean ± SEM from three biological replicates. Statistical significance was determined by one-way ANOVA followed by Tukey’s multiple comparisons test (*P* < 0.1).

Following rifampin treatment, two antioxidant pathway metabolites showed trends toward altered abundance in Δ*pstA1* compared to wild-type: ergothioneine and 5-oxoproline (p < 0.1, ANOVA; Fig. 7B). Ergothioneine, a sulfur-containing antioxidant, showed a trend toward elevation in Δ*pstA1* compared to both wild-type and complement strains. This trend is notable given that *whiB3*, which was upregulated 2.6-fold in the transcriptomic analysis and confirmed by RT-qPCR, is a metabolic regulator involved in the production of ergothioneine and mycothiol, which play an important role in detoxifying the oxidative stress [35]. Similarly, 5-oxoproline, a glutamate derived metabolite, showed a trend toward increased abundance in the deletion mutant [36]. The observed trends suggest that Δ*pstA1* may exhibit hypersensitivity to rifampin largely because of alterations in antioxidant metabolism following increased rifampin exposure.

## Discussion

Components of the Pst system have been implicated in virulence, stress tolerance, and antibiotic susceptibility in several bacterial pathogens, suggesting that phosphate-specific transport pathways may represent promising targets for therapeutic intervention [14, 37]. This study establishes PstA1 as a critical determinant of rifampin tolerance in *Mtb* through mechanisms distinct from drug resistance. Deletion of *pstA1* resulted in enhanced rifampin killing without substantial change in MIC, a phenotype that remained consistent across nutrient starvation conditions and correlated with reduced survival in IFN-γ-activated macrophages. Integrative multi-omics analysis revealed that *pstA1* loss triggers constitutive phosphate starvation signaling, leading to PE/PPE porin upregulation. These findings raise the possibility that PE/PPE porin upregulation enhances membrane permeability, thereby promoting rifampin uptake. Metabolic profiling revealed nonsignificant upregulation of antioxidant pathways including ergothioneine and glutathione biosynthesis, suggesting elevated oxidative stress resulting from increased intracellular rifampin accumulation due to hyperpermeability [35].

In contrast to prior studies showing that *pstA1* deficiency reduces the MIC of rifampin by 8-fold, we found a more modest 2-fold decrease in the rifampin MIC against Δ*pstA1*. This discrepancy may be due to differences in the parent strain used (Erdman vs. H37Rv) [20]. Notwithstanding these findings, we observed significantly increased rifampin activity against Δ*pstA1* in time-kill assays, the standard approach for assessing antibiotic tolerance [5].

Δ*pstA1* retained its rifampin hypersusceptibility under nutrient starvation and showed reduced survival under phosphate depletion, consistent with previous reports [14, 15] and validating its core metabolic function in phosphate acquisition. This metabolic vulnerability extended to host-pathogen interactions. While a previous Tn-seq study found that *pstS3*, *pstC2*, *pstA1*, and *phoT* were each important for macrophage survival under all activation conditions tested [15], Δ*pstA1* showed a more nuanced phenotype in the current study, with no survival defect in resting macrophages but significant reduction in intracellular survival in IFN-γ-activated macrophages by day 4. This finding aligns well with that from a separate study reporting that Δ*pstA1* retained full virulence in IFN-γ⁻/⁻ mice but showed reduced chronic-phase survival in wild-type mice, establishing that PstA1 is specifically required for surviving IFN-γ-mediated immunity [14].

To understand the mechanistic basis for enhanced rifampin susceptibility, we examined drug uptake dynamics and cellular structure. We observed accelerated rifampin uptake with significantly elevated drug accumulation by 6 hours, implicating increased drug permeability and/or reduced efflux. Electron microscopy revealed no major structural changes in cellular morphology. Our transcriptomic analysis provided insight into the molecular basis for this altered permeability state. In the absence of rifampin exposure, Δ*pstA1* showed extensive cellular reprogramming with 58 genes differentially expressed. Most striking was the constitutive activation of the phosphate starvation regulon, including *pstS3*, *pstC2*, *senX3*, and *regX3,* indicating that cells lacking functional PstA1 exist in a state of perceived phosphate limitation regardless of environmental phosphate availability. The marked upregulation of *pstS3* and *pstC2* likely represents compensatory transcriptional upregulation consistent with constitutively activated RegX3, as multiple studies have demonstrated RegX3-dependent regulation of the *pstS3-pstC2-pstA1* operon [13, 14, 18]. Without the functional PstA1 permease, however, this transcriptional response cannot restore phosphate transport, resulting in persistent phosphate starvation signaling.

This baseline stress state likely compromises bacterial ability to respond effectively to additional challenges like antibiotic exposure. There is substantial concordance between our transcriptomic data and those of another study evaluating a *pstA1* deletion mutant in the Erdman background, as 22 common DEGs were also identified in both studies, including *senX3*, *regX3*, the *pstS3-pstC2-pstA1* operon, *whiB3*, and 9 PE/PPE genes, validating the core transcriptional signature of *pstA1* deficiency across strain backgrounds [14].

Among the 36 genes differentially expressed in *ΔpstA1* regardless of rifampin treatment, 12 encoded PE/PPE family genes and 4 encoded ESAT-6-like proteins. Four of these PE/PPE genes (PE18, PPE19, PPE25, PPE27) are part of the *esx-5* locus, consistent with established regulation of ESX-5 secretion by the Pst/SenX3-RegX3 axis and the finding that *pstA1* deletion causes ESX-5 substrate hypersecretion. This upregulation provides a direct mechanistic link to the increased rifampin uptake we observed [16]. Recent work demonstrated that Δ*pstA1* mutants overcome acid growth arrest by upregulating PE19-PPE51 porin expression to enhance glycerol uptake [12], while another study established that PE/PPE proteins function as substrate-specific outer membrane channels [38]. The constitutive overexpression of multiple PE/PPE proteins in Δ*pstA1* facilitates not only glycerol uptake but also rifampin entry through increased membrane permeability, potentially explaining the accelerated drug accumulation and enhanced killing [12]. Transcriptomic analysis revealed no downregulation of known rifampin efflux pump genes, further supporting the hypothesis that enhanced drug accumulation results from increased membrane permeability rather than impaired efflux. The absence of gross structural defects on electron microscopy supports the interpretation that increased permeability results from specific changes in channel composition rather than general membrane disruption.

Interestingly, DNA-interacting genes were only upregulated in Δ*pstA1* in the absence of rifampin, suggesting that drug’s inhibitory effects on RNA polymerase may suppress some baseline transcriptional differences. We identified only a few genes differentially regulated exclusively upon rifampin exposure, likely because of the relatively low drug concentration and brief timepoints, which were selected to minimize global transcriptional inhibition.

Our integrated metabolomic and transcriptomic analysis revealed activation of antioxidant pathways in rifampin-treated Δ*pstA1*, consistent with elevated oxidative stress from increased intracellular drug accumulation. We observed an upward trend of metabolites involved in antioxidant defense, including 5-oxoproline, glutamate, glutathionylspermidine, and notably ergothioneine. The ergothioneine response is particularly informative: this mycobacterial antioxidant provides protection against rifampin-induced damage, as a study demonstrated that Δ*egtD* mutants lacking ergothioneine show enhanced susceptibility to rifampin and other first-line drugs [39]. The apparent paradox in Δ*pstA1*, where ergothioneine is elevated yet susceptibility increases, could reflect compensatory antioxidant upregulation that is insufficient to overcome the primary defect of PE/PPE-mediated drug hyperpermeability.

The concurrent 2.6-fold upregulation of *whiB3*, a redox-responsive regulator of thiol metabolism, further supports activation of oxidative stress defenses. While one study showed that WhiB3 negatively regulates ergothioneine biosynthesis under standard conditions, the simultaneous increase in both *whiB3* expression and ergothioneine levels in rifampin-treated Δ*pstA1* suggests that rifampin-induced oxidative stress may trigger compensatory antioxidant production [35]. The elevation of 5-oxoproline, a glutamate-derived metabolite, indicates broad activation of thiol-based defenses beyond ergothioneine alone [40]. Critically, despite these activated protective pathways, Δ*pstA1* exhibits marked hypersusceptibility to rifampin, suggesting that the primary driver of enhanced killing is increased drug accumulation rather than deficient antioxidant capacity.

This is further supported by recent findings showing that WhiB3 expression is induced by both acidic pH and phosphate limitation through RegX3-dependent mechanisms, and that elevated WhiB3 levels in Δ*pstA1* correlate with reduced acid-induced ROS and preserved GAPDH activity [12].

While increased rifampin uptake provides mechanistic connection to enhanced killing, the molecular basis for altered drug accumulation requires further investigation. The link between PE/PPE upregulation, ESX-5 hypersecretion, and rifampin permeability needs direct demonstration. The transcriptomic analysis captures a snapshot of cellular responses and may miss dynamic changes occurring at other timepoints. The targeted metabolomics, while revealing specific antioxidant pathway alterations, can be extended to look at relevant perturbations in central metabolism or lipid composition that could contribute to the phenotype.

The identification of *pstA1* as a tolerance determinant opens potential therapeutic avenues, and targeting this pathway requires careful consideration. Our findings show that loss of PstA1 function creates multiple vulnerabilities, including enhanced drug uptake, metabolic dysregulation, and impaired stress responses, suggesting that constitutive activation of phosphate-sensing pathways during antibiotic treatment becomes a liability rather than an adaptive advantage. The specific mechanisms identified are intriguing targets: the WhiB3-ergothioneine stress response axis and PE/PPE-mediated membrane changes represent downstream effectors that could be modulated without disrupting fundamental phosphate homeostasis. Understanding how phosphate transport influences antibiotic tolerance could inform adjunctive strategies to eliminate persister populations, potentially shortening the lengthy treatment regimens currently required for TB cure.

## Materials and Methods

### Bacterial Strains and Plasmids

*Mycobacterium tuberculosis* H37Rv served as the wild-type strain for all experiments. Gene deletion strains were constructed using ORBIT (Oligonucleotide-Recombineering followed by Bxb1 Integrase Targeting) cloning as previously described [94]. Wild-type H37Rv was first transformed with pKM444, a non-integrating plasmid encoding RecT recombinase and Bxb1 integrase (gift from Kenan Murphy, University of Massachusetts). In a second electroporation, cells received pKM464 (also from Kenan Murphy) along with a gene-specific oligonucleotide designed to delete pstA1. This oligonucleotide contained the first and last 11 codons of pstA1 flanked by 37-bp homology arms, with the Bxb1 attP integration site positioned centrally. Transformants were selected on antibiotic-selective plates, and deletion junctions were verified by PCR amplification followed by Sanger sequencing.

For complementation, the pstA1 coding sequence plus its upstream regulatory region (422 bp upstream of pstS3 plus the first 75 bp of pstS3) was PCR-amplified from wild-type H37Rv genomic DNA. This fragment was cloned into pMH94, an episomal mycobacterial vector modified with apramycin resistance (gift from Dr. Gyanu Lamichhane, Johns Hopkins University), using In-Fusion Snap Assembly (Takara Bio). The vector was first linearized by restriction digestion with EcoRI and XbaI, gel-purified using QIAquick Gel Extraction Kit (Qiagen), then assembled with the PCR product. Cloned constructs were transformed into Stellar Competent Cells (Takara Bio), selected on LB plates with appropriate antibiotics, and verified by PCR before electroporation into the ΔpstA1 strain.

### Culture Conditions and Media

*M. tuberculosis* strains were routinely cultured in Middlebrook 7H9 broth supplemented with 10% OADC enrichment, 0.2% glycerol, and 0.05% Tween-80, or on Middlebrook 7H11 agar plates incubated at 37°C. Phosphate-depleted medium was prepared as described previously [24] by omitting inorganic phosphate salts from the 7H9 formulation. THP-1 monocytic cells were maintained in RPMI-1640 medium (Gibco) containing 2 mM L-glutamine and 10% heat-inactivated fetal bovine serum at 37°C with 5% CO₂.

### Operon Structure Validation

To confirm co-expression of *pstS3*, *pstC2*, and *pstA1* as a single operon, RNA was extracted from logarithmically growing *Mtb* H37Rv cultures using TRIzol reagent. Following DNase-I treatment to eliminate genomic DNA contamination, RNA was reverse transcribed using SuperScript VILO cDNA Synthesis Kit (Invitrogen). PCR primers were designed to amplify both intragenic regions (within individual genes) and intergenic regions (spanning gene boundaries).

### Whole Genome Sequencing Validation

High-molecular-weight genomic DNA was extracted from 10 mL log-phase cultures using an extended protocol optimized for mycobacteria. Cultures were lysed by sequential treatments with lysozyme (20 hours at 37°C), SDS/proteinase K (50°C), and CTAB, followed by chloroform/isoamyl alcohol extraction and isopropanol precipitation. To verify deletion and complementation, genomic DNA from wild type, Δ*pstA1*, and complement strains were submitted to SeqCenter (Pittsburgh, PA) for whole genome sequencing. Libraries were prepared using the Illumina DNA Prep kit with tagmentation-based fragmentation and sequenced on an Illumina NovaSeq X Plus platform (2×151 bp paired-end reads). Raw sequencing reads were aligned to the *M. tuberculosis* H37Rv reference genome, and coverage analysis was performed using SAMtools v1.18. Deletion boundaries were confirmed by examining read coverage across the *pstA1* locus.

### Drug Susceptibility Testing

Minimum inhibitory concentrations (MICs) were determined by broth microdilution in 96-well plates. Bacterial suspensions were prepared by washing log-phase cultures and resuspending in Tween-80-free 7H9 medium to OD₆₀₀ 0.01. Equal volumes (100 μL) of bacterial suspension and 2× serial drug dilutions were combined and incubated at 37°C for 6 days. Metabolic activity was assessed by adding 50 μL alamarBlue reagent (1:1 mixture of 10× alamarBlue and 10% Tween-80) and measuring fluorescence after 24 hours (544/590 nm excitation/emission) on a Fluostar OPTIMA plate reader (BMG Labtech).

### Transmission Electron Microscopy

Transmission electron microscopy was performed previously [27]. Briefly, mid-log phase bacteria were collected and resuspended in 3% formaldehyde, 2.0% glutaraldehyde, 80 mM Sorenson’s phosphate, and 4 mM MgCl2, pH 7.2 with duplicate samples for negative staining and sectioning. Samples were fixed overnight at 4°C before processing and imaging. Cell wall thickness was measured at two locations per bacterium: one in a thin region and one in a thick region when present, to account for regional variations.

### Time-Kill Assays

For nutrient-rich conditions, bacterial cultures were grown to mid-logarithmic phase in complete 7H9 medium, then diluted to OD₆₀₀ 0.05 one day prior to drug exposure. On day 0, rifampin was added to a final concentration of 2 μg/mL unless otherwise noted (stock: 2 mg/mL in DMSO), with vehicle controls receiving equivalent DMSO volumes. Each condition was performed in triplicate using 5 mL cultures in 50 mL conical tubes, incubated at 37°C with shaking. Bacterial viability was quantified by serial dilution plating on 7H11 agar at days 2, 4, and 6 post-treatment. Plates were incubated at 37°C for 14-21 days before colony counting.

For nutrient starvation conditions, log-phase cultures were washed and resuspended in PBS containing 0.05% Tween-80 to OD₆₀₀ 0.5, then incubated without shaking at 37°C for 14 days to establish nutrient-deprivation. Rifampin (2 μg/mL) or vehicle control was then added, and bacterial survival was monitored by CFU plating as described above.

For phosphate depletion experiments, cultures were washed twice and resuspended in phosphate-free 7H9 medium to OD₆₀₀ 0.05. Growth and survival were monitored over time by CFU enumeration, with samples taken at indicated intervals during incubation at 37°C with shaking.

### Macrophage Infection Assays

THP-1 cells were differentiated by treatment with 50 ng/mL phorbol 12-myristate 13-acetate (PMA) for 48 hours at 37°C with 5% CO₂. Differentiated macrophages were optionally activated with 20 ng/mL recombinant IFN-γ for 24 hours before infection. Bacterial suspensions were prepared from log-phase cultures and used to infect macrophages at a multiplicity of infection (MOI) of 1:1. After 4 hours, extracellular bacteria were removed by washing three times with PBS, and fresh medium was added. At specified time points, macrophages were lysed with 0.05% SDS and lysates were plated on 7H11 agar for CFU determination.

### Polyphosphate Quantification

Polyphosphate levels were measured as described previously [41]. Bacterial pellets were lysed in guanidine thiocyanate buffer (5 M guanidine thiocyanate, 50 mM Tris-HCl pH 7.0) using bead beating (3 cycles, 20 seconds at 6,800 rpm). After centrifugation, supernatants were stored at −20°C, and protein concentrations were determined using DC Protein Assay (Bio-Rad) for normalization. Polyphosphate was isolated using the GeneClean III Kit (MP Biomedicals) following manufacturer protocols, including RNase/DNase treatment to remove nucleic acids. Polyphosphate concentrations were determined fluorometrically using DAPI staining (100 μg/mL final concentration) with excitation/emission at 415/525 nm.

### Transcriptomic Analysis

Wild-type and Δ*pstA1* cultures were grown to mid-log phase (OD₆₀₀ ∼0.5) in 45 mL 7H9 medium. To capture early transcriptional responses, cultures were treated with subinhibitory rifampin (7.5 ng/mL, 1/4 MIC) or vehicle. RNA samples were collected at baseline, 30-or 60-minutes post-treatment with three independent biological replicates per condition.

Bacterial pellets (15 mL culture) were resuspended in TRIzol with 0.1 mm zirconia beads and lysed using a Precellys Evolution homogenizer (3 cycles, 30 seconds at 7,400 rpm).

RNA was extracted with chloroform, precipitated with ethanol, and purified on RNeasy columns (Qiagen) with on-column DNase I digestion. Following polyA selection, strand-specific libraries were prepared using NEBNext Ultra II Directional RNA Library Prep Kit and sequenced on Illumina NovaSeq X Plus (2×150 bp paired-end) at the Johns Hopkins Genetic Resources Core Facility.

Raw reads were quality-assessed (FastQC v0.12.1) [30] and aligned to the *M. tuberculosis* H37Rv reference genome (Bowtie2 v2.5.1) [31]. Gene counts were quantified with featureCounts (Subread v2.0.0) [32]. Differential expression analysis used DESeq2 v1.44.0 with significance thresholds of adjusted p-value < 0.05 (Benjamini-Hochberg) and |log₂FC| > 1. Pathway analysis was performed using KEGGREST to query KEGG pathways [34] for significantly differentially expressed genes.

### RT-qPCR Validation

Twelve genes were analyzed by RT-qPCR, including eleven target genes and sigA as a housekeeping control (18-22 nt, 100-147 bp amplicons) designed with IDT PrimerQuest (Supplementary Table S1). Total RNA (500 ng) was DNase-treated, purified on RNeasy columns, and reverse transcribed with iScript RT Master Mix (Bio-Rad) alongside no-RT controls. qPCR used 10 μL reactions with SYBR Green Supermix on a QuantStudio 3 system (95°C/10 min; 40 cycles: 95°C/15s, 60°C/1 min). Relative expression was calculated using the ΔΔCt method with sigA normalization.

### Metabolite extraction and LC-MS analysis

*Mtb*-laden filters were incubated on m7H10 plates at 37°C for 5 days to mid-log phase [35]. Cells were quenched with prechilled acetonitrile:methanol:H₂O (40:40:20, v:v:v) at −40°C. Metabolites were extracted through mechanical lysis using 0.1-mm zirconia beads in a Precellys tissue homogenizer for 2 min at 6,000 rpm, repeated three times under continuous cooling at ≤2°C. Cell lysates were clarified by filtration through 0.22-μm Spin-X columns. Residual protein content was measured with a BCA protein assay kit (Thermo Scientific) to normalize metabolite ion counts to cell biomass.

Extracted metabolites were separated using a Cogent Diamond Hydride Type C column with mobile phases of solvent A (ddH₂O with 0.2% formic acid) and solvent B (acetonitrile with 0.2% formic acid). An Agilent 6230 TOF mass spectrometer was coupled to an Agilent 1290 LC system. Dynamic mass axis calibration was maintained via continuous infusion of reference mass solution through an isocratic pump with a 100:1 splitter. This setup achieved mass errors of ∼5 ppm and mass resolution of 10,000-25,000 over m/z range 62–966, with a dynamic range of 5 log₁₀. Ions were identified based on unique accurate mass and retention time. Data analysis used Agilent Profinder B.07.00 with mass tolerance <0.005 Da. All metabolomics data represent the average of at least two independent triplicates.

### Antibiotic permeability (rifampin uptake)

Rifampin uptake was measured using a previously validated device that enables timed start-stop measurements of antibiotics in spent media [35, 36]. The device comprises a plastic inset containing m7H9 with known antibiotic concentration. *Mtb*-laden filters were generated using wild-type, ΔpstA1, and complement strains and placed onto media containing 1 μg/mL rifampin at 37°C. After incubating for 2, 6, and 24 hours, bacterial lysates and spent m7H9 were analyzed for rifampin content by LC-MS. Rifampin uptake was calculated as: [RIF]drug only – [RIF]filtrate, with three biological replicates.

### Statistical Analysis

Data are presented as mean ± standard deviation unless otherwise noted. Statistical comparisons were performed using appropriate tests as specified in figure legends (unpaired t-tests, one-way or two-way ANOVA with post-hoc corrections). Multiple comparisons were corrected using the Holm-Sidak method or Tukey’s Multiple Comparisons where applicable. Statistical significance was defined as p < 0.05. Sample sizes are indicated in individual figure legends.

## Code and Data Availability

Code to replicate the performed analysis is available at https://doi.org/10.5281/zenodo.17344250. Raw sequencing data has been deposited to the Gene Expression Omnibus (GEO) and can be accessed at GSE310692. Additional data supporting the findings of this study are available upon request.

## Acknowledgements

We thank Dr. Kenan Murphy (University of Massachusetts) for generously providing the pKM444 and pKM464 plasmids used for ORBIT cloning, and Dr. Gyanu Lamichhane (Johns Hopkins University) for providing the pMH94 episomal vector. We are grateful to Megan Gelement (Johns Hopkins University) for contributions to methodology and data analysis discussions. We are grateful to SeqCenter (Pittsburgh, PA) for whole genome sequencing services. We thank the Johns Hopkins Genetic Resources Core Facility (GRCF) for RNA sequencing services.

## Author Contributions

Conceptualization: CD, PCK; Methodology: CD, GVAC, JSB, GYL, HE, PCK; Data acquisition: CD, GVAC, GYL; Data analysis: CD, GVAC, GYL; Supervision: JSB, HE, PCK; Funding acquisition: PCK, HE; Writing of original draft: CD, GVAC, PCK; Writing— review & editing: CD, GVAC, JSB, GYL, HE, PCK

## Supplementary Figures

**Figure S1.**
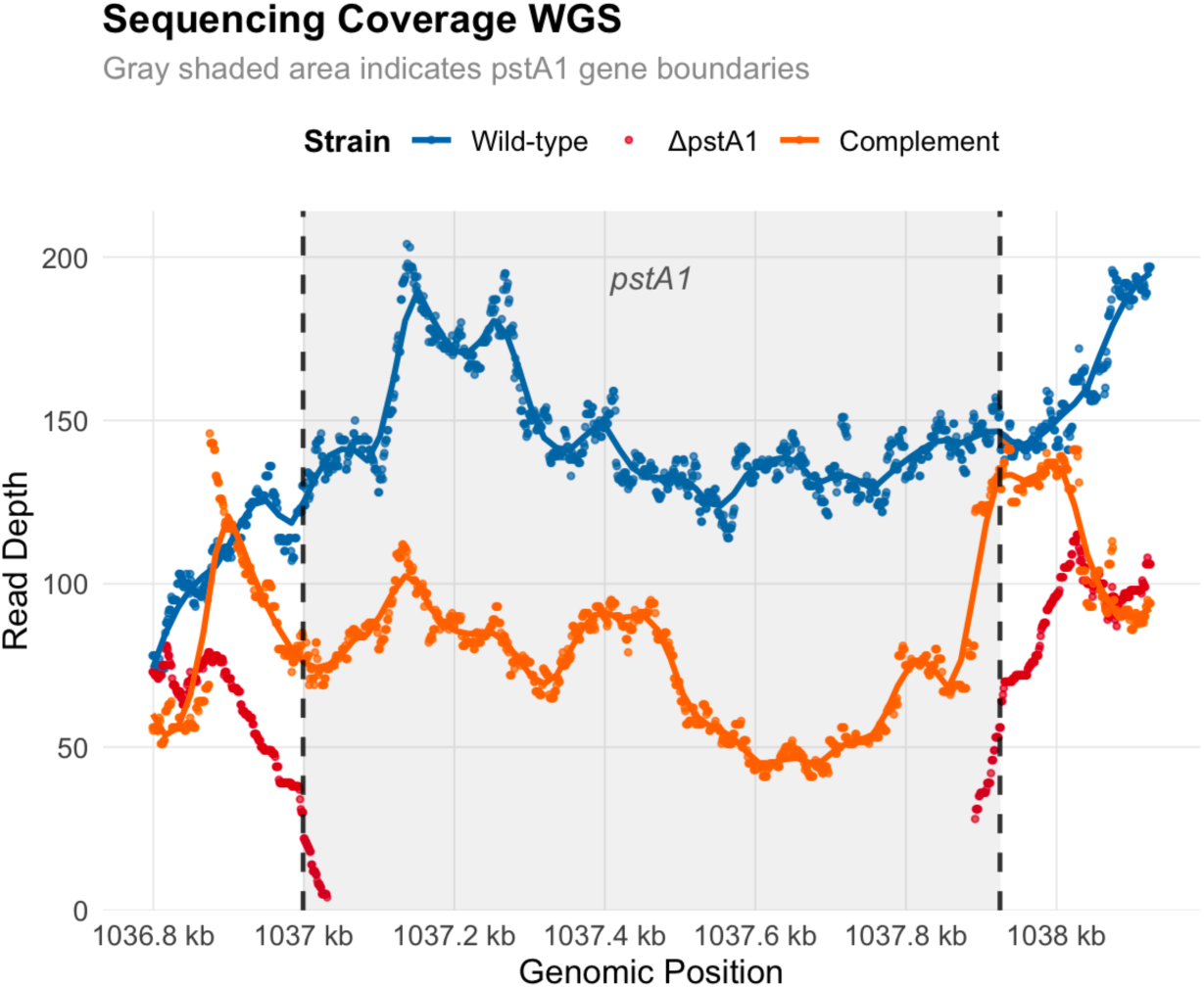
Whole genome sequencing of pstA1 deletion and complementation strains. Read depth coverage across the *pstA1* locus in wild-type H37Rv (blue), ΔpstA1 deletion mutant (red), and Δ*pstA1::pstA1* complement strain (orange). Gray shaded region indicates pstA1 gene boundaries (coordinates 1,036,999-1,037,925 bp). Vertical dashed lines mark gene start and end positions. Wild-type exhibits uniform coverage (∼150-200× read depth) throughout the *pstA1* coding sequence. The Δ*pstA1* strain shows near-zero coverage within the deleted region with maintained coverage in flanking sequences, confirming successful targeted deletion. The complement strain displays reduced coverage (∼50-100× read depth) within *pstA1* relative to wild-type, consistent with episomal expression from the pMH94 plasmid. Individual points represent sequencing depth at each nucleotide position; smoothed trend lines (LOESS, span = 0.1) are shown for wild-type and *pstA1* complement strains. Coverage was calculated from reads aligned to the *M. tuberculosis* H37Rv reference genome using samtools depth and visualized in R.

**Figure S2.**
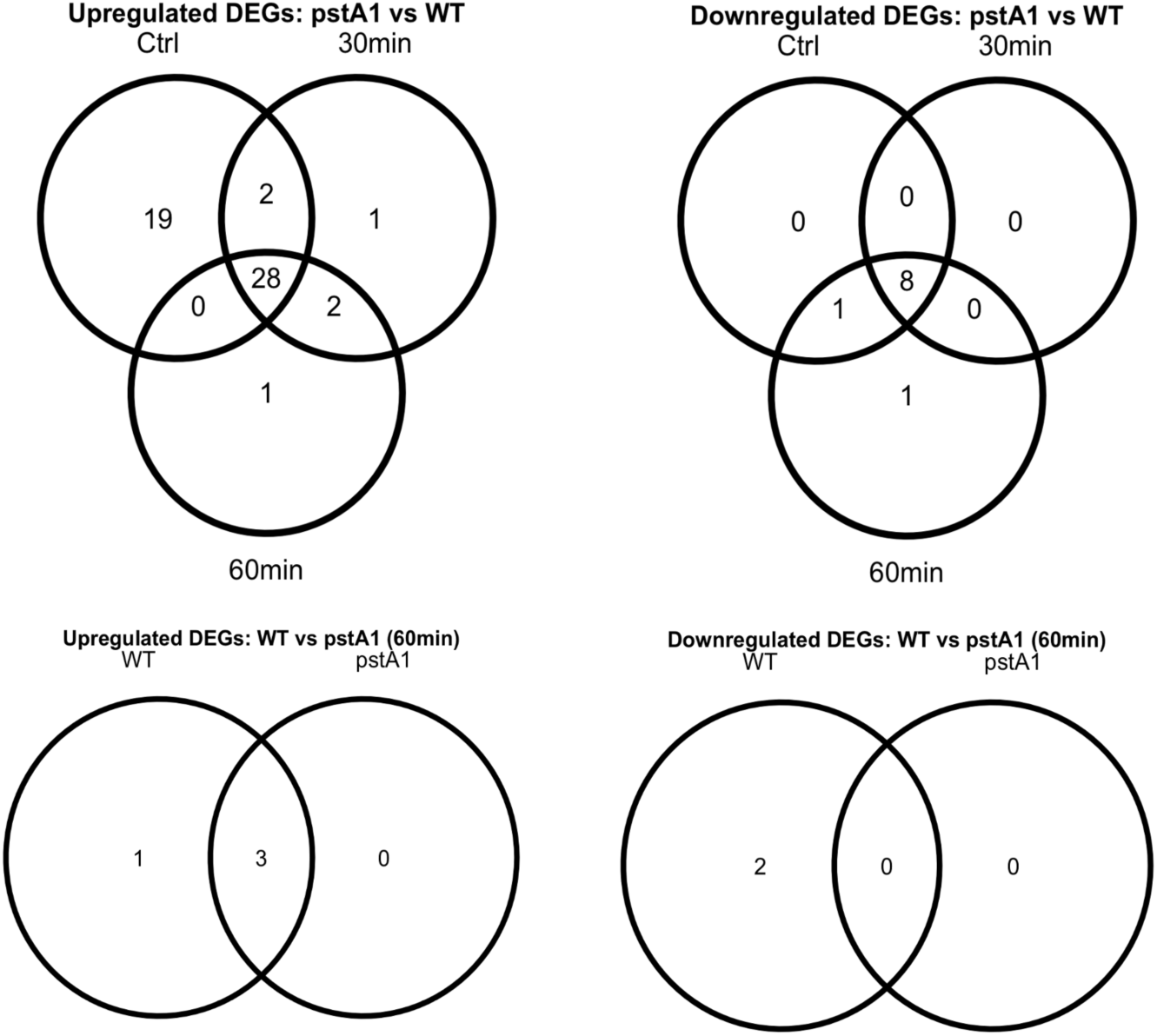
Venn diagram analysis of differentially expressed genes between Δ*pstA1* and wild-type strains. Overlap of differentially expressed genes (DEGs) identified by RNA-seq (log2 fold change > 1, adjusted p-value < 0.05). Top panels show upregulated (left) and downregulated (right) genes in Δ*pstA1* compared to WT across control, 30 min, and 60 min rifampin treatment conditions. Bottom panels show upregulated (left) and downregulated (right) genes comparing WT and Δ*pstA1* responses to 60 min rifampin treatment relative to their respective untreated controls. Numbers indicate the count of unique or shared DEGs in each category.

**Figure S3.**
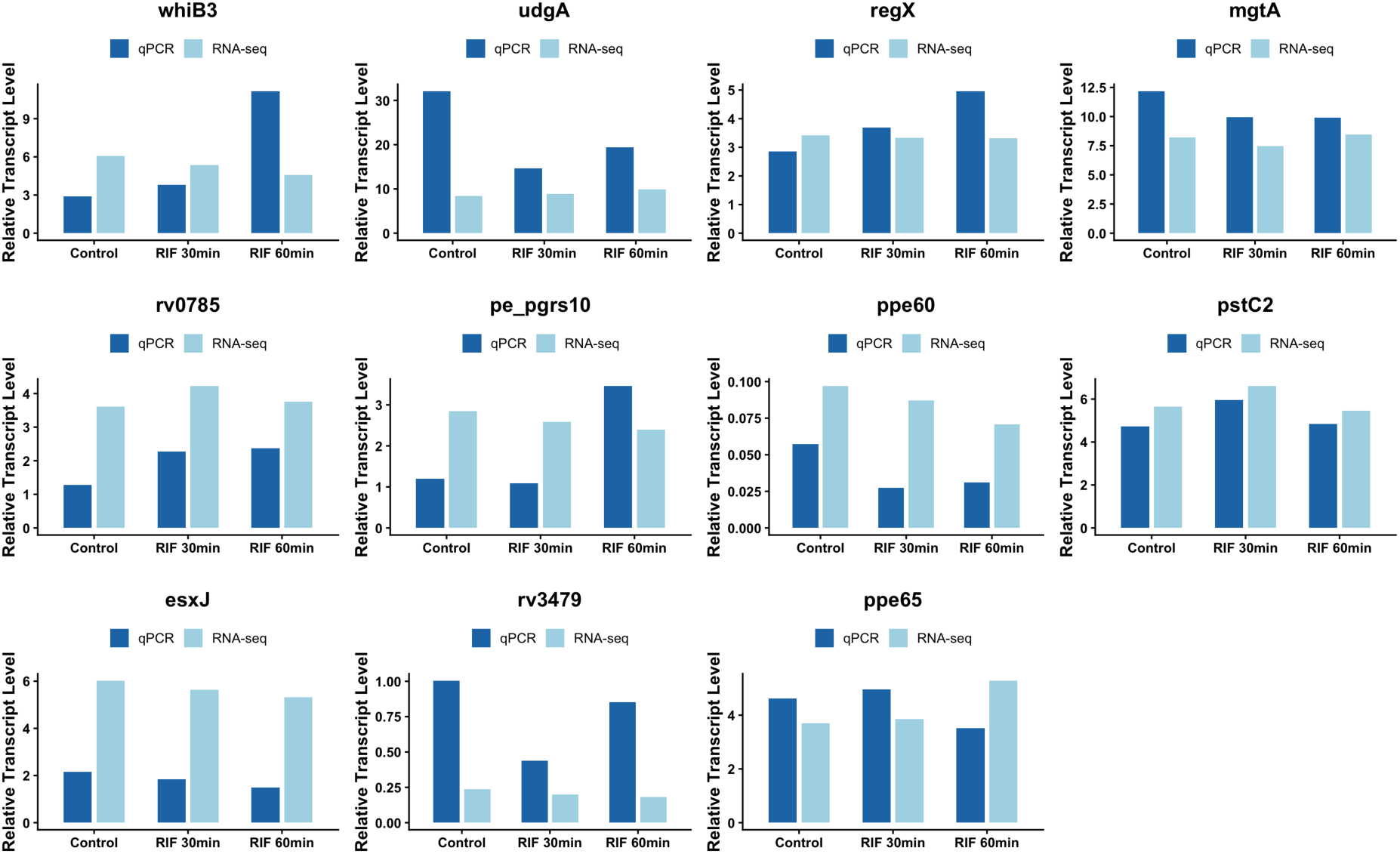
Validation of RNA-seq results by quantitative PCR. Comparison of transcript levels measured by RNA-seq (light blue) and qPCR (dark blue) for selected differentially expressed genes in wild-type *M. tuberculosis* under control conditions and following 30- or 60-min rifampin treatment. Genes were selected to represent diverse functional categories including transcriptional regulators (whiB3, regX), metabolic enzymes (udgA, mgtA), PE/PPE family proteins (pe_pgrs10, ppe60, ppe65, rv3479), phosphate transport (pstC2), and ESX secretion (esxJ). Relative transcript levels are normalized to control conditions. Strong correlation between RNA-seq and qPCR measurements validates the RNA-seq dataset.

**Supplementary Table S1.**
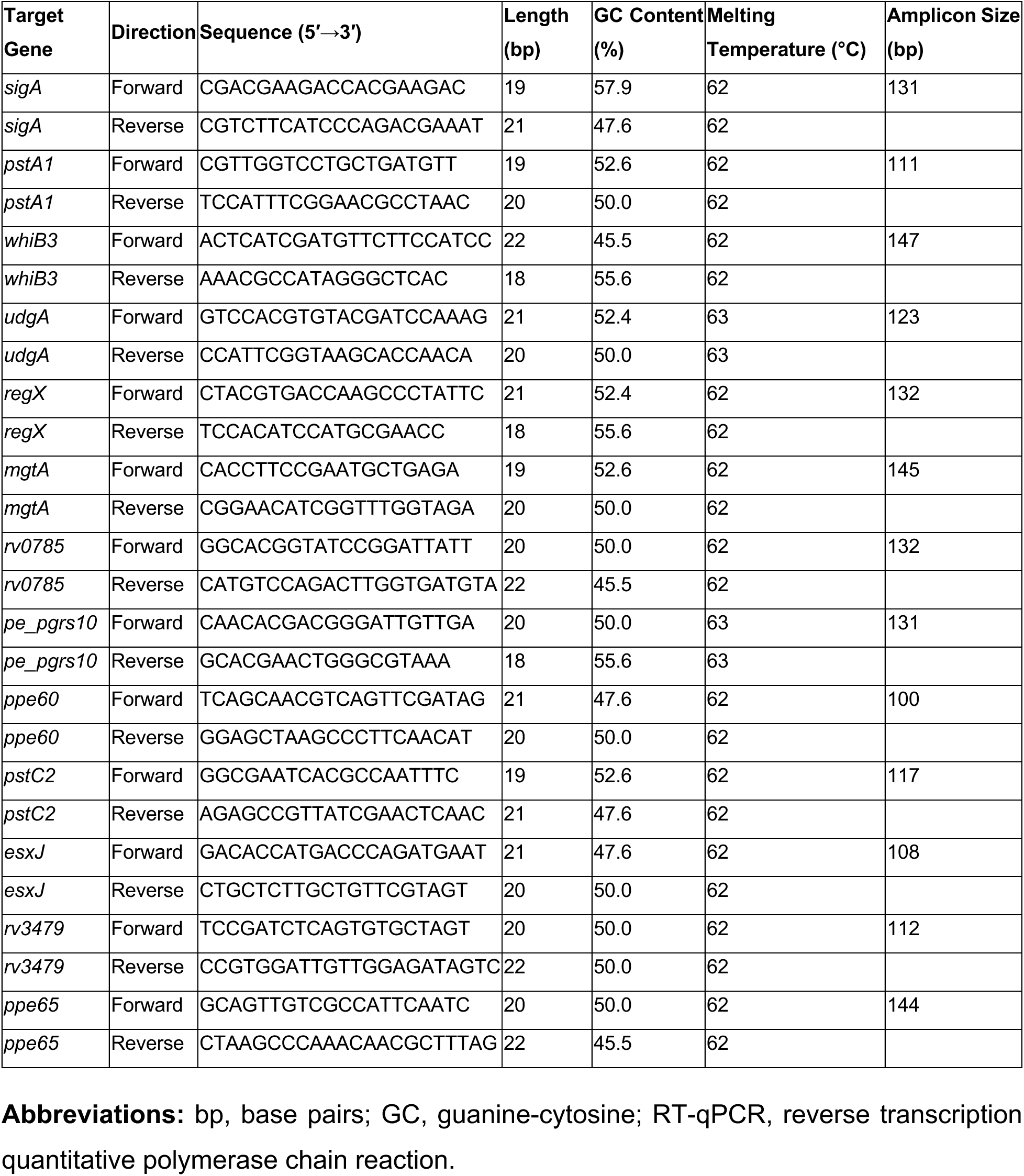
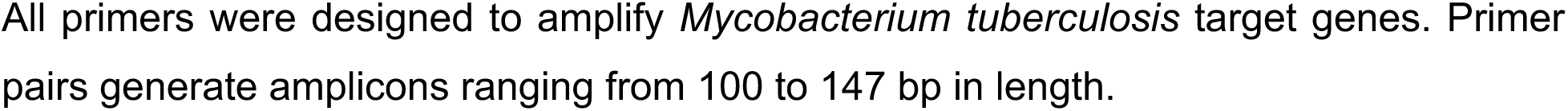
Oligonucleotide primers used for RT-qPCR analysis of *Mycobacterium tuberculosis* genes.

**Supplementary Table S2.** All genes meeting the significance threshold (|log2 fold change| > 1, adjusted p-value < 0.05) across seven pairwise comparisons are listed. Comparisons include temporal responses to rifampin treatment in wild-type and Δ*pstA1* strains (30 and 60 min versus untreated controls), and strain-specific differences between Δ*pstA1* and wild-type under control and rifampin-treated conditions. For each gene, the following information is provided: comparison name, *M. tuberculosis* H37Rv gene identifier (Rv number), gene name, log2 fold change, adjusted p-value (Benjamini-Hochberg correction), and KEGG pathway annotations where available. A total of 58 genes were differentially expressed between Δ*pstA1* and wild-type under control conditions (49 upregulated, 9 downregulated), 41 at 30 min rifampin (33 upregulated, 8 downregulated), and 41 at 60 min rifampin (31 upregulated, 10 downregulated). Only 6 genes responded to rifampin in wild-type (4 upregulated, 2 downregulated at 60 min), and 3 genes in Δ*pstA1* (all upregulated at 60 min), highlighting the substantial transcriptional differences between the strains.

